# The microbial community dynamics of cocaine sensitization in two behaviorally divergent strains of collaborative cross mice

**DOI:** 10.1101/2022.12.06.519360

**Authors:** Thi Dong Binh Tran, Christian Monroy Hernandez, Hoan Nguyen, Susan Wright, Center for Systems Neurogenetics of Addiction, Lisa M. Tarantino, Elissa J. Chesler, George M. Weinstock, Yanjiao Zhou, Jason A. Bubier

**Author notes:** Correspondence: Jason A. Bubier, The Jackson Laboratory, 600 Main Street, Bar Harbor ME 04609.

## Abstract

The gut-brain axis is increasingly recognized as an important pathway involved in cocaine use disorder. Microbial products of the murine gut have been shown to affect striatal gene expression, and depletion of the microbiome by antibiotic treatment alters cocaine-induced behavioral sensitization in C57BL/6J male mice. Some reports suggest that cocaine-induced behavioral sensitization is correlated with drug self-administration behavior in mice. Here we profile the composition of the naïve microbiome and its response to cocaine sensitization in two Collaborative Cross (CC) strains. These strains display extremely divergent behavioral responses to cocaine sensitization. A high-responding strain, CC004/TauUncJ (CC04), has a gut microbiome that contains a greater amount of *Lactobacillus* than the cocaine-nonresponsive strain CC041/TauUncJ (CC41). The gut microbiome of CC41 is characterized by an abundance of *Eisenbergella, Robinsonella* and *Ruminococcus*. In response to cocaine, CC04 has an increased *Barnsiella* population, while the gut microbiome of CC41 displays no significant changes. PICRUSt functional analysis of the functional potential of the gut microbiome in CC04 shows a significant number of potential gut-brain modules altered after exposure to cocaine, specifically those encoding for tryptophan synthesis, glutamine metabolism, and menaquinone synthesis (vitamin K2). Depletion of the microbiome by antibiotic treatment revealed an altered cocaine-sensitization response following antibiotics in female CC04 mice. Depleting the microbiome by antibiotic treatment in males revealed increased infusions for CC04 during a cocaine intravenous self-administration dose-response curve. Together these data suggest that genetic differences in cocaine-related behaviors may involve the microbiome.

**FUNDING:** U01DA043809 to JAB, GMW P50DA039841 to EJC

## 1. INTRODUCTION

In the face of the COVID-19 pandemic, the substance use disorder (SUD) and overdose epidemic has continued to grow [1, 2]. Overdose deaths spiked at the start of the pandemic and stayed high throughout 2020 [3]. Among drugs of abuse, cocaine use disorder is highly heritable, with ∼70% of the variation ascribed to genetics in twin studies [4]. Advanced genetically diverse mouse strains such as Collaborative Cross (CC) are well suited for exploring the genetic contribution to complex diseases such as cocaine use disorder [5, 6]. Within this population of mice are 45 million SNPs segregating, a level of genetic diversity comparable to that of the human population. The CC mouse population is a panel of recombinant inbred (RI) mice derived from an eight-way cross of distinct sequenced inbred mouse strains. All mice within a CC strain are homozygous and the CC itself is inbred and represents a reproducible population. Approximately 60 CC strains are commercially available. In addition to their diverse phenotypes and host genetics, the microbiome of these recombinant inbred lines exhibits tremendous variation [7, 8]. When controlled for diet and environment, host genetics explains a large part of the variation in microbiome composition [9]. Thus, host genetics may control cocaine use disorder by influencing the microbiome.

The role of the microbiome in SUD is gaining increasing recognition (reviewed in [10, 11]). Previous studies in mice have shown that antibiotic ablation of the microbiome alters low-dose cocaine sensitization and conditioned place preference (CPP) in male C57BL/6J mice [12]. The enhanced CPP response from antibiotics was corrected by short-chain fatty acid administration, suggesting that an intact microbiome ‘represses’ an endogenous cocaine response. A similar reversible response was also reported for sensitization and CPP in response to morphine in male C57BL/6 mice. [13] Recently treatment of male C57BL/6J mice with Proteobacteria was shown to alter cocaine neurobehavioral responses in a glycine dependent manner [14]. Finally, work in the alcohol field using the binge drinking model, drinking in the dark, using C57BL/6J mice given a 2-week antibiotic pretreatment showed significantly increased alcohol consumption [15]; which this increase could be reversed by sodium butyrate supplementation [16]. However opposite results were obtained in a Wistar derived high drinking selected rat strain in which antibiotic ablation reduced their consumption of ethanol[17]. Ezquer, F., et al restored the high drink phenotype with oral administration of *Lactobacillus rhamnosus*, suggesting the intact microbiome encourages alcohol consumption[17]. Collectively these studies suggest a role for the microbiome in substance use disorder and related mouse behaviors.

We examined two previously reported behaviorally divergent CC strains, CC004/TauUnc (CC04) and CC041/TauUnc (CC41) [18]. These strains exhibit extreme genetic and biological variation underlying resistance or vulnerability to the stimulatory and reinforcing effects of cocaine. CC04 mice exhibit a robust locomotor response to cocaine and demonstrate active operant cocaine intravenous self-administration. Conversely, CC41 are non-responsive to the locomotor stimulatory effects of cocaine and are do not acquire cocaine intravenous self-administration under the same conditions [18]. To determine whether the microbiome contributes to observed cocaine behavior in these CC04 and CC41, we profiled the fecal microbiome before mice entered a cocaine sensitization protocol and 19 days later on completion of the protocol by 16S rRNA gene sequencing. We identified distinct microbiome composition and microbiome metabolic pathways in naïve CC04 and CC41 strains, and showed a number of gut-brain modules were altered in the CC04 strain in response to cocaine. Finally, we sought to investigate causation using an antibiotic ablation intervention followed by both cocaine sensitization and cocaine intravenous self-administration in the same strains. In total. we demonstrate that the microbiomes of the CC04 and CC41 strains are unique to each strain and differentially responsive to cocaine. The impact of this work is a demonstration that the role of the microbiome in affecting behavioral response to cocaine is strain, sex and dosage specific.

## 2. MATERIALS AND METHODS

### 2.1 Animals

The CC004/TauUncJ and CC041/TauUncJ are CC RI strains obtained from The Jackson Laboratory. They were bred in an elevated barrier Pathogen & Opportunistic-Free Animal Room (https://www.jax.org/-/media/jaxweb/health-reports/ax12.pdf?la=en&hash=970E1B297070DD9022BB3CCA38A195F02B4ED295) and transferred at ween to an intermediate barrier specific pathogen-free room (https://www.jax.org/-/media/jaxweb/health-reports/g3b.pdf?la=en&hash=914216EE4F44ADC1585F1EF219CC7F631F881773). All animal protocols were reviewed and approved by The Jackson Laboratory Institutional Animal Care and Use Committee (Approval #10007).

All mice were group-housed with same-sex siblings until six weeks of age in duplex, individually vented cages (Thoren Caging Systems, Inc. Pennsylvania, USA; Cage #3) with pine-shaving bedding (Hancock Lumber) and environmental enrichment consisting of a nestlet and a Shepherd Shack^®^ (Shepard Specialty Papers). Mice were singly housed at six weeks and began behavioral phenotyping at eight weeks of age. Mice were maintained on a 12:12 light cycle (lights on at 6 AM) and allowed *ad libitum* access to the standard rodent chow [sterilized NIH31 5K52 6% fat chow (LabDiet/PMI Nutrition, St. Louis)] and acidified water (pH 2.5–3.0) supplemented with vitamin K. The mice were identified by ear notching at weaning and moved between cages by forceps. Cages were changed once a week. If a cage change was scheduled for a testing day, a clean cage was prepared, and post-test, mice were returned to a new, clean cage. The Jackson Laboratory follows husbandry practices following the recommendations of the American Association for the Accreditation of Laboratory Animal Care (AAALAC).

### 2.2 Phenotyping

#### 2.2.1 Open Field Apparatus

The open field apparatus was a square-shaped, clear polycarbonate arena (Med-Associates #MED-OFAS-515U) with dimensions 17.5 inches length x 17.5 inches width x10.0 inches height (44.5 cm × 44.5 cm × 25.4 cm). External to the arena’s perimeter at the floor level, on the left and right sides, is a pair of horizontal infrared photo beam sensors (16 × 16 beam array). An additional pair of infrared photo beam sensors raised 3 inches from the arena floor (16 × 16 array) are situated at the arena’s front and rear outer sides and used to capture vertical activity. Each arena is placed within a sound-attenuating, ventilated cabinet with interior dimensions: 26”W x 20”H x 22”D (Med Associates, #MED-OFA-017) containing a fan that provides white noise. Each cabinet contains two incandescent lights, each affixed in the upper rear two corners of the cabinet at the height of approximately 18.5 inches from the center of the arena floor which provides illumination of 60±10 lux when measured in the center of the arena floor. Data was collected using Activity Monitor software version 7.0.5.10 (SOF-812; Med Associates, Inc; RRID: SCR_014296).

#### 2.2.2 Cocaine Sensitization

On all test days, mice were transported from the housing room on a wheeled rack and left undisturbed to acclimate in an anteroom adjacent to the procedure room for a minimum of 30 minutes. Before the first mouse was placed into any behavioral apparatus and between each subject, the equipment was thoroughly sanitized with 70% ethanol solution (in water), and wiped dry with clean paper towels. After all testing for the day, the subjects were returned to the housing room and the arenas were sanitized with Virkon (Lanxess), followed by 70% ethanol to remove any residue. Two different sensitization paradigms were used.

##### 2.2.2.1 Cocaine Sensitization using 10 mg/kg

Complete phenotyping details can be found on the Mouse Phenome Database (https://phenome.jax.org/projects/CSNA03/protocol?method=open+field+test). The cocaine sensitization paradigm was carried out as described in [18]. Briefly, cohorts of mice (3 males, 3 females) of each strain were assigned to sham or cocaine treatment cohorts. Mice were tested for 90 minutes each test day in a 19-day protocol. On days 1, 2 and 12, all animals received saline; on days 3, 5, 7, 8, 11 and 19, half the animals received cocaine (10 mg/kg), while the other half received saline. On each test day, mice were placed in the arena for 30 minutes, then injected with saline or cocaine and returned to the arena for 60 minutes. Before the start of the sensitization paradigm, fresh fecal boli were collected from each mouse. Following the 19-day sensitization paradigm, fecal pellets were collected 60 minutes after the last cocaine injection. Fecal samples were processed and analyzed as described below.

##### 2.2.2.2 Cocaine Sensitization using 5 mg/kg

Following antibiotic treatment (described below), a different cocaine sensitization paradigm was utilized to replicate the protocol of Kiraly 2016 [12]. Briefly, mice were given saline on days one, two and three, and on days 4, 5, 6, 7 and 8, mice were given cocaine at 5 mg/kg i.p. Immediately following cocaine or saline administration, mice were placed in the open field and behavior was monitored for 45 minutes. Numerous behaviors were recorded in the open field as described in Table S1. Before the start of the sensitization paradigm, fresh fecal boli were collected from each mouse. At the completion of testing fecal boli were collected. Fecal samples were processed and analyzed as described below.

### 2.3 Antibiotic Treatment

Fecal boli samples were collected from treatment-naïve mice. At 8-10 weeks of age, mice started antibiotic treatment (ABX). Mice were administered sulfatrim (19.75 mg/liter sulfamethoxazole + 3.95 mg/liter trimethoprim) and 1 g/L of ampicillin sodium salts, 1g/L of Metronidazole, 0.5 g/L Vancomycin hydrochloride, and 1 g/l of Neomycin (all pharmaceutical grade) in their drinking water continuously. Sweetener (2.5 g/liter, aspartame) was added to the antibiotic water and the water of a second cohort of control (CTRL) mice [7, 19]. One week following the initiation of antibiotic treatment, which was continuous throughout testing, mice from both cohorts were tested in the open field, light-dark and holeboard. Following this baseline testing, 48 mice of both sexes and strains underwent cocaine sensitization in two batches of 24. After testing, fecal boli were harvested from both groups, and cecal contents and striatum were dissected.

### 2.4 Fecal Collection

Mice were placed in a clean cage for five minutes. Any fecal pellets deposited were collected, placed in Eppendorf tubes, and stored at -80 C. If no pellets were produced in the first five minutes, mice were left in the cage for a longer period.

### 2.5 Dissections

To control for circadian effects, behavioral testing and euthanasia were consistently performed between 8-12 AM. All surgical instruments were cleaned with RNAase away between each animal (ThermoFischer Scientific). The mice were euthanized by decapitation rather than cervical dislocation in order to maintain the brain structure. The cecum was identified, and the fecal contents were extruded into an Eppendorf tube and flash-frozen on dry ice. All snap-frozen samples were stored at −80°.

### 2.6 16S Microbiome Analysis

DNA was isolated using the Shoreline complete V1V3 kit (Shoreline Biome, cat #SCV13) per the manufacturer’s instructions and used to create barcoded V1V3 amplicons. Briefly, ∼5mg of mouse fecal pellet was lysed with a combination of heat, pH, and cell wall disruptors in a single step. DNA was recovered using the magnetic beads, and an aliquot from each sample was transferred to a well in the provided PCR plate containing sample barcode/Illumina sequencing primers. Next, 2x PCR mix from the kit was added, and PCR was performed according to the manufacturer’s instructions. After PCR, samples were pooled, purified using a MinElute PCR Purification Kit (Qiagen, cat# 28004) and diluted for sequencing on the Illumina MiSeq (RRID: SCR_020134) platform generating 2×300 paired reads. Sequencing reads are processed by removing the sequences with low quality (average qual <35) and ambiguous codons (N’s). Paired amplicon reads are assembled using Flash. Chimeric amplicons were removed using UChime software (RRID: SCR_008057) (73). Our automatic pipeline used the processed reads for operational taxonomic unit (OTU) generation. Each OTU will be classified from phylum to genus level using the most updated RDP classifier and training set (RRID: SCR_006633). A taxonomic abundance table was generated with each row as bacterial taxonomic classification, each column as sample ID and each field with taxonomic abundance. The abundance of a given taxon in a sample was presented as relative abundance (the read counts from a given taxon divided by total reads in the sample).” We also used PICRUSt2 (RRID: SCR_022647) [20] to predict the functional profiling of the bacterial communities by ancestral state reconstruction using 16S rRNA gene sequences. Following the protocol described by Valles-Colomer [21], gut-brain module analysis was performed on the PICRUST2 results. Multiple testing correction of the Gut-Brain Modules was performed using the qvalue package (RRID: SCR_001073).

### 2.7 Metagenomic whole genome shotgun (mWGS) sequencing, Data Processing and Metagenomics Species Detection

Cecal feces from both cocaine sensitization experiments (with and without antibiotics) were analyzed by mWGS. Libraries are constructed with an average insert size of 500 bases and then sequenced on the HiSeq2500 instrument producing 150 base read pairs from each fragment, yielding ∼3 million read pairs/sample. Following mWGS sequencing, sequence data was run through a quality control pipeline to remove poor-quality reads and sequencing artifacts. Sequencing adapters were first removed using TRIMMOMATIC (RRID: SCR_011848) [22]. Next, exact duplicates, low quality, low complexity reads and mouse DNA contamination were removed using GATK-Pathseq (RRID: SCR_005203)[23] pipeline with a k-mer-based approach. For optimization of mouse DNA decontamination, we have built a new GATK-Pathseq 31-mer database by concatenating the following collection of DNA sequences: MM10_GRCm38 reference; sixteen diverse laboratory mouse reference genomes define strain-specific haplotypes and novel functional loci; NCBI UniVec clone vector sequences; repetitive element sequences from RepBase23.02 database; and mouse encode transcripts databases (v25). The final clean reads were used for taxonomic classification and metabolic function analysis for further downstream analysis.

An optimized GATK-Pathseq classification pipeline is time efficient and robust solution for taxonomic classification at the species level. This pipeline used BWA-MEM alignment (minimum 50 bp length at 95% identity). It mapped the final clean reads to the latest updated reference of microbial genomes built by concatenating RefSeq Release 99 (March 2nd, 2020) nucleotide FASTA sequence files of bacteria, viruses, archaea, fungi, and protozoa. Gatk-Pathseq All read counts of microbial species of all kingdoms are used for species abundance analysis.

### 2.8 Differential abundance analysis of microbial genes and metabolic pathways

The KEGG ortholog (KO) profiling was performed by HumanN2 (RRID: SCR_016280)[24]. Using the DESeq2 package (RRID: SCR_015687)[25] dedicated to performing comparative metagenomics, the inference of the abundance of genes and pathways was obtained and visualized using a volcano plot. Because of the potential high false positive rate of DESeq [26], we plotted raw and relative abundance to inspect the results. Following the protocol described by Valles-Colomer [21], gut-brain module analysis was performed on the mWGS results. Multiple testing correction of the Gut-Brain Modules was performed using the qvalue package (RRID: SCR_001073).

### 2.9 Biogenic amines analysis

#### 2.9.1 Tissue Extraction

Tissues were frozen at -80º C and shipped to the Vanderbilt Neurochemistry Core for analysis. The samples were held on dry ice prior to the addition of homogenization buffer to prevent the degradation of biogenic amines. Tissues were homogenized using a handheld sonic tissue dismembrator in 100-750 ul of 0.1M TCA containing 0.01M sodium acetate, 0.1mM EDTA, and 10.5 % methanol (pH 3.8). Ten microliters of homogenate were used for the protein assay. The samples were spun in a microcentrifuge at 10,000 g for 20 minutes. Then, the supernatant was removed for HPLC-ECD analysis. HPLC was performed using a Kinetix 2.6um C18 column (4.6 × 100 mm, Phenomenex, Torrance, CA, USA). The same buffer used for tissue homogenization is used as the HPLC mobile phase. The following biogenic amines were analyzed Norepinephrine (NE), 3,4-Dihydroxyphenylacetic acid (DOPAC), dopamine (DA), 5-hydroxyindoleacetic acid (5-HIAA), homovanillic acid (HVA), serotonin (5-HT),3-methoxytyramine (3-MT).

#### 2.9.2 Protein assay

Protein concentration in cell pellets was determined by BCA Protein Assay Kit (Thermo Scientific). Ten microliter tissue homogenate is distributed into a 96-well plate and 200 µl) mixed BCA reagent (25 ml of Protein Reagent A is mixed with 500 μl of Protein Reagent B) is added. Incubate the plate at room temperature for two hours for color development. A BSA standard curve is run at the same time. Absorbance was measured by the plate reader (POLARstar Omega), purchased from BMG LABTECH Company.

### 2.10 Cocaine Intravenous self-administration

Mice were tested following a modification of the protocol described in Dickson *et al*. [27]. Mice >12 weeks of age underwent indwelling jugular vein catheterization under oxygen/isoflurane anesthesia by The Jackson Laboratory surgical services. The catheter (Instech, C20PU-MJV, Plymouth Meeting, PA) was routed through the stainless coupler of a mesh button. The mesh button was sutured in place subcutaneously. The port was flushed with saline and filled with 10 ul of lock solution (Lumen lock solution; 500 IU heparin/ml lock solution). The catheter was capped with a protective aluminum cap (Instech., PHM-VAB95CAP). The mouse was observed for at least three days after surgery to ensure the incision was healing properly. If mice exhibited pain signs, buprenorphine (0.05mg/kg SQ) was administered every 4-5 hours or carprofen (5 mg/kg SQ) was administered every 24 hours. Mice were provided a minimum ten-day post-operative recovery period in their home cage before testing. Mice were transported in their home cages from the housing room to the procedure room on a wheeled transport rack. Before the start of testing, upon transport to the procedure room, mice were weighed (once a week) and briefly handled to be assessed for any welfare concerns that may result in exclusion from testing (e.g., wounds or looking sickly). IVSA data were collected using Med Associates operant conditioning chambers (307W), fitted with two retractable levers on the front wall flanking the right and left sides of the center panel food hopper. Directly over each lever (∼ 2-3 inches above) were red stimulus lights. A house light (ENV-315W) was centrally mounted on the rear wall. A modified Plexiglas floor, fabricated at the Jackson Laboratory, was fitted to cover the metal floor grids. The chambers were within sound attenuating cubicles (ENV-022MD). A 25-gauge single-channel stainless steel swivel was mounted to a counterbalanced lever arm attached to the outside of the chamber. Tubing was used to connect a syringe mounted on the infusion pump to the swivel and to connect the swivel to a vascular access harness. Operant conditioning chambers were controlled by a Med Associates control unit using MED-PC IV software (RRID:SCR_012156).

Mice began cocaine IVSA testing on a fixed-ratio 1 (FR1) schedule at a dose of 1.0 mg/kg/infusion of cocaine hydrochloride (NIDA Drug Supply). Infusions from the active lever were at 8.85 ul/sec, with 1 sec per 10g mouse weight. Each IVSA session was two hours each day. After each day of testing, Baytril (enrofloxacin) was administered by catheter to the mice at 22.7 mg/kg, followed by a 20 ul injection of Heparin lock solution (100 U/ml heparin/saline). Acquisition criteria were met when mice had 5 (could be non-consecutive) sessions with ≥10 infusions. The mouse reached stabilization when the number of infusions didn’t vary more than 20% for the last two consecutive sessions. Mice were moved onto the dose response curve even if the had not reached acquisition or stabilization criteria at 18 days. Following stabilization, the mice were tested on 1.0 mg/kg through a dose-response curve from high to low doses. Doses were presented in the following order: 1.0, 0.32, 0.1, 0.032 mg/kg/infusion. Mice were tested on consecutive days on the same dose until stabilization criteria were met or a max of 5 sessions, then moved on to the next dose. Mice may fail to survive the surgery, have a non-patent catheter, a systemic infection, a local infection at the catheter port, cocaine overdose, or air embolism. These mice were excluded from the study. Final mouse numbers in IVSA were CC04 antibiotic treatment n=5, control aspartame water n=4, CC41 antibiotic treatment n=5, control aspartame water n=7.

### 2.11 Statistics

Microbial community analysis was performed by R version 3.5.1 (RRID: SCR_001905). Principal coordinate analysis (PCoA) plots, boxplots and heatmaps were generated for graphical visualization using Phyloseq, ggplot2 (RRID: SCR_014601)[28] and ComplexHeatmap (RRID: SCR_017270)[29] packages. Richness was calculated as the number of OTUs present in each sample. The Shannon Diversity Index combined species richness, and the evenness was computed as Σp_i_*ln(p_i_), where pi presents the proportional abundance of species. The non-parametric Wilcoxon or Kruskal-Wallis rank sum-tests were used for differential diversity or abundance between two or more groups and corrected for multiple comparisons by the Benjamini-Hochberg procedure. Beta diversity was analyzed at the OTU level using the Bray-Curtis distance for community abundance and the Jaccard distance for community presence/absence.

The among-group differences were determined using the permutational multivariate analysis of variance by the distance matrices (ADONIS). These tests compare the intragroup distances to the intergroup distances in a permutation scheme and then calculate a p-value. These functions are implemented in the Vegan package (RRID: SCR_011950)[30]. For all permutation tests, we used 10,000 permutations. Statistical analyses for sensitization and neuropeptides (sensitization only) were conducted using JMP 16 (SAS Institute; RRID: SCR_014242). The best model is

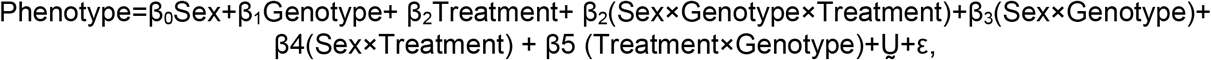

where ε is random error. The parameters (β, ε) were estimated by Type III non-sequential ordinary least squares in the ANOVA model. Repeated measures analysis of variance was performed using MANOVA to estimate the effects of strain, sex, treatment and their interaction. Ṵ represents the symmetric nature of the variance co-variance matrix of the random effects. In all cases, the full model was fit and reduced by dropping non-significant inter-actions followed by main effects. In order to determine the nature of differences detected in the ANOVA model, planned contrasts were performed giving the terms not included in the model a weight of zero and giving the terms to be compared (treatment, strain) values of opposite weights, −1 and +1.

## 3. Results

### 3.1 Fecal microbiome composition of CC004 and CC041 lines

We sought to determine the fecal microbiome composition of two strains of CC mice shown to be behaviorally divergent in response to cocaine[18]. We performed 16S V1-V3 sequencing on fecal boli from naïve male and female mice and mice that had completed a 19-day cocaine sensitization paradigm. The behavioral response of the strains was as previously published [18] and our results replicated those (**Figure S1)**. A principal coordinates analysis of the beta-diversity of the gut microbiome showed significant separation of the microbiomes by strain (PC1 35.8%) and cocaine effect (PC2 13.4%) (**Figure 1A**). Analysis of Group Dissimilarity-Adonis based on microbial abundance matrix with Bray-Curtis distance showed a significant difference between the naïve and cocaine exposure groups in the CC04, [Adonis F_(3,22)_ = 4.3768 p <0.001, R2=0.40866]. This cocaine effect was strain specific [CC04 F_(1,10)_ =2.6019, p<0.05 R2=0.186; CC41 F_(1,11)_ =1.2826, p=0.17 R^2^=0.011]

**Figure 1.**
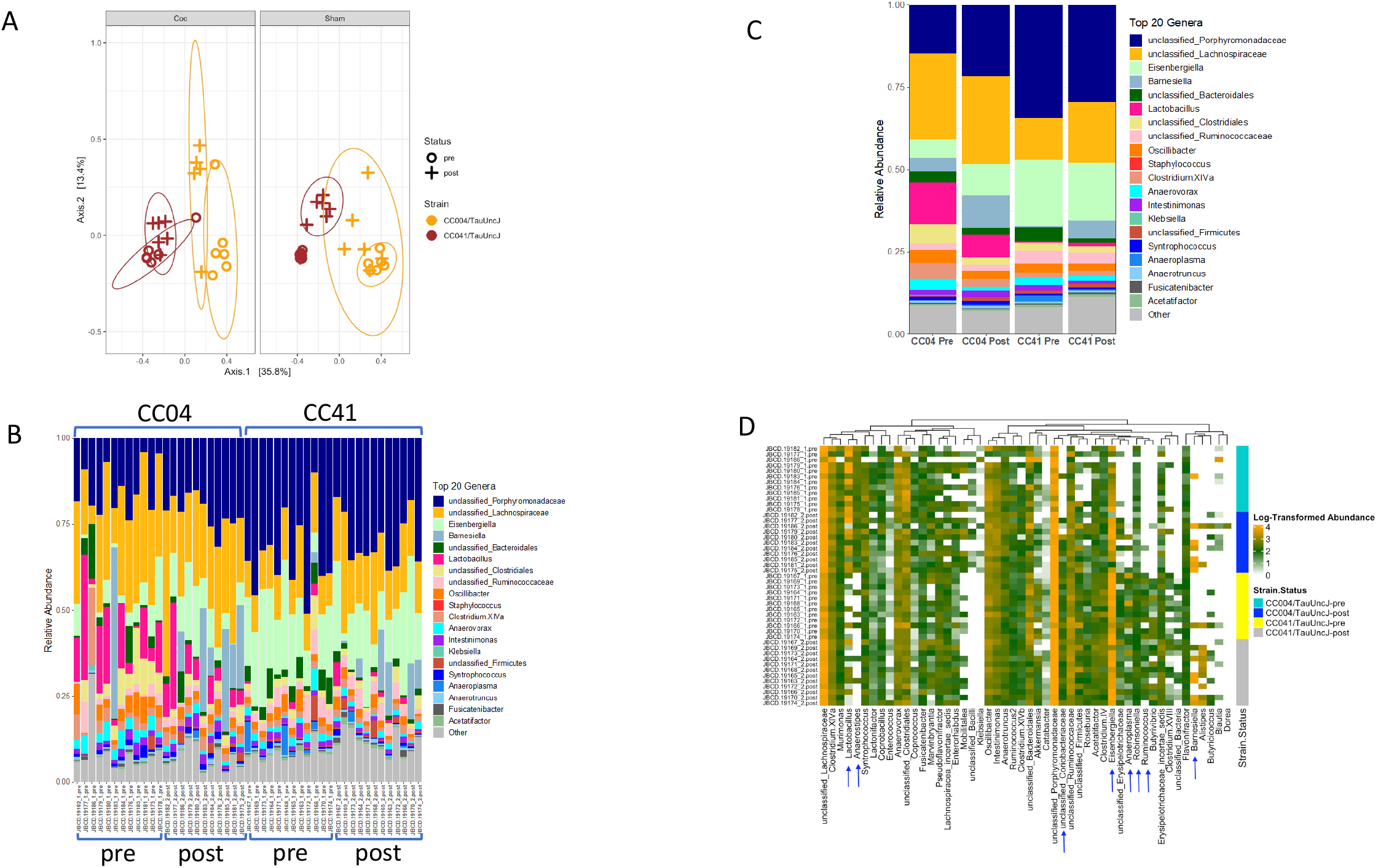
Microbial composition of two naïve Collaborative Cross strains and in response to cocaine sensitization paradigm. **A**. Principal components analysis of bacterial beta diversity at OTU level using Bray-Curtis for 16S microbiome data showing clustering of samples by host genotype (PC1 35.8%) and Treatment (PC2 13.4%). Three males and three females of each strain were tested with cocaine or a sham treatment. **B**. The percent abundance of the 20 most abundant genera in female and male mice of strain CC04 and CC041 before a cocaine sensitization paradigm and after a cocaine sensitization paradigm. **C**. Strain and treatment average microbiome composition of the top 20 genera. **D**. Heat map representation of the microbiome composition of CC04 and CC041 pre and post cocaine sensitization. White color corresponds to absence of bacteria while light green to orange colors correspond to low and high abundance of bacteria. Arrows show important bacteria that significantly differentiate between strain and cocaine sensitization.

Naïve CC04 displayed decreased *Eisenbergella* (W=9, Z=-3.80, q<0.0001 r=0.79), *Aneroplasma* (W=0, Z=-4.27, q<0.001, r=0.89), *Robinsonella* (W=23 Z=-2.69 q=0.018, r=0.56), and *Ruminococcus* (W=5.5, Z=-3.98, q<0.001, r=0.83) and an increase in *Lactobacillus* (W=126, Z=-3.69, q=0.001, r=0.77), and *Anaerostipes* (W=128, Z=-3.90, q <0.0001, r=0.813), compared to Naive CC41. (**Figure 1B, Figure 1D** with arrows). In response to cocaine, CC04 showed an increase in *Barnsiella* (W=0, Z=-3.63, q=0.0082, r=0.93) and *unclassified_Coriobacteriaceae* (W=0, Z=-3.60, q<0.0082, r=0.93). CC41 showed no significant differences that passed a stringent FDR=0.05 after cocaine treatment. *Barnsiella* did show an increase (W=20.5, Z=-1.92, p=0.05, q=0.34, r=0.45) in CC41 but it did not pass multiple correction testing. A full table of differences, (p<0.05 and q>0.05) between strains can be found in **Table S2**.

### 3.2 PICRUSt analysis of the 16S data identifies molecular functions that differ between strains and those that are affected by cocaine treatment in CC04 mice

Based upon the linkage between phylogeny and function, Phylogenetic Investigation of Communities by Reconstruction of Unobserved States (PICRUSt) is an approach to predictive metagenomics that has been demonstrated to provide useful insights into thousands of uncultivated microbial communities [31]. PICRUSt was used to compare the functional properties of the microbiome before the sensitization paradigm with the microbiome after the sensitization paradigm in both CC04 and CC41 mice. Outlier samples (19183 CC04 pre and 19179 CC04 Post) were removed for this analysis. These samples appeared to be switched due to their position in the PCA, although results were similar when they were included. There were 50 Gut-Brain KEGG Modules that were significantly different, after correcting for multiple testing, between naïve CC04 and naïve CC41. (**Table S3, Figure 2A,B**). The KO0686 other glutamine showed greatest upregulation (4.48 of log2 fold change) in CC04 mice, compared to CC41 mice. log fold change (LFC) of 4.48, and the most downregulated was KO12942 other_glutamate LFC -6.08. Other modules represented multiple times included Menaquinone synthesis (vitamin K2), Isovaleric acid synthesis I (KADH pathway), acetate and propionate metabolism. The KEGG ontology pathways, represented by CC04 microbiome post-sensitization, included 29 Gut-Brain KEGG Modules that were significantly different after correcting for multiple testing **(Table S3, Figure 2C,D**). There were three downregulated modules related to tryptophan synthesis, four downregulated modeules related to glutamine metabolism, seven modules related to Menaquinone synthesis (vitamin K2), and others related to glutamate and GABA synthesis. The greatest fold change was -7.06 LFC for other_glutamine K12942 (q=0.007).

**Figure 2.**
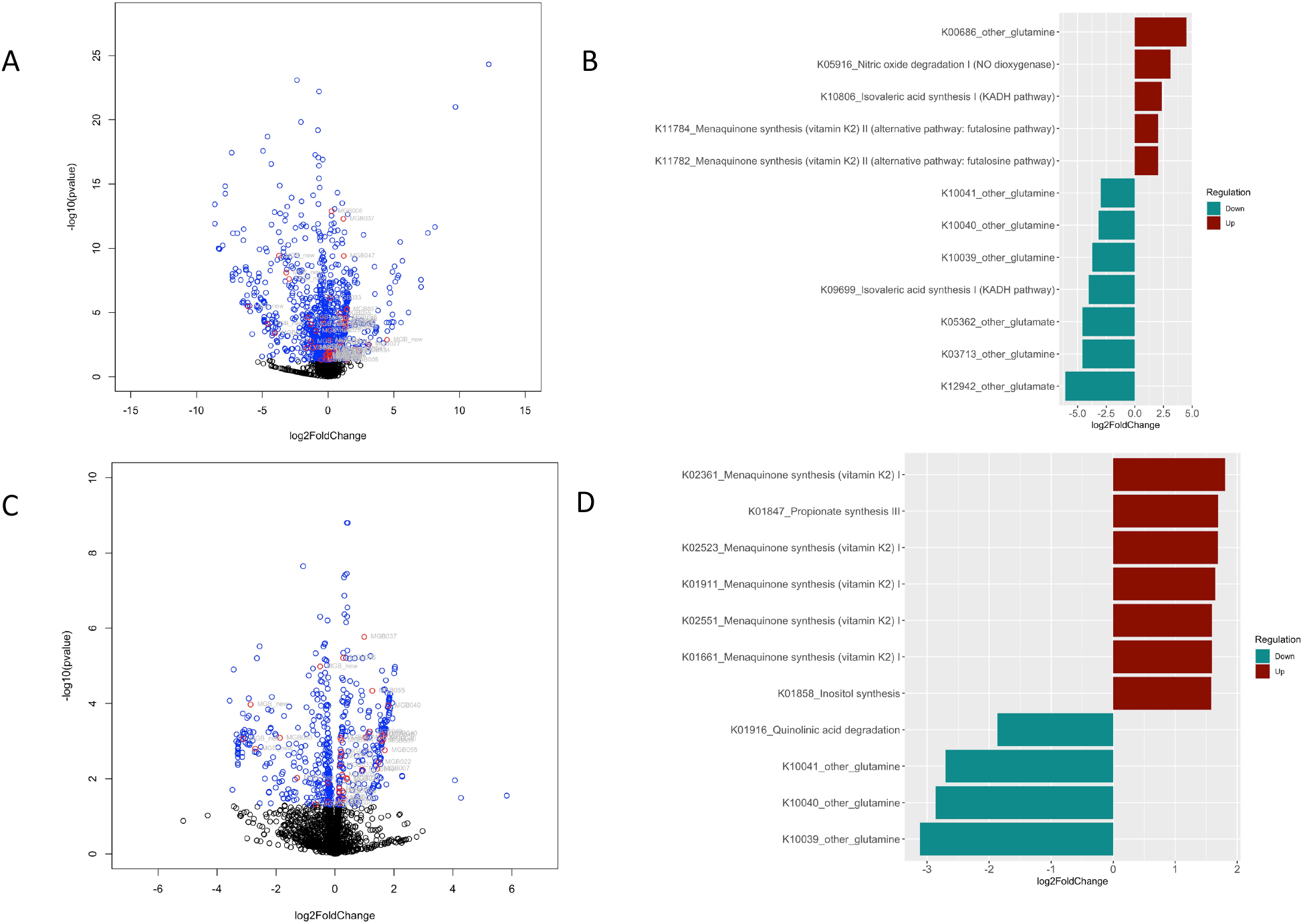
Volcano plots of the functional categories encoded by the microbiome. **A**. A scatterplot showing the statistical significance and magnitude of change of the KO clusters as determined by PICRUSt2 of 16S data between CC04 and CC041 pre sensitization. Blue= q >0.05, Red= q<0.05. **B**. The gut-brain module KO categories (q <0.05) from PICRUSt2 of 16S data. **C**. A scatterplot showing the statistical significance and **D**. magnitude of change of the gut-brain module KO clusters as determined by PICRUSt2 of 16S data between CC04 pre sensitization and CC04 post sensitization. **\***Outlier samples 19183 CC04 pre and 19179 CC04 Post were removed from the later PICRUSt analysis.

### 3.3 Antibiotic treatment effects on cocaine sensitization behavior

Previous work has shown the involvement of the microbiome in cocaine sensitization and CPP in male C57BL/6J mice at low doses of cocaine (5 mg/kg) [12]. CC04 and CC41 mice were given *ad libitum* access to antibiotic-treated water or control (aspartame) water starting at 8-10 weeks of age (**Figure 3**) to test for the involvement of the microbiome in observed differences in cocaine sensitization response in these two strains. The antibiotic treatment didn’t cause any adverse effect on the mice as measured by weight differences between the control and treated groups (**Figure S2**). In a repeated-measures ANOVA of the full factorial model that was sequentially reduced there were no significant strain × sex × treatment effects or interactions and only a univariate effect of sex was observed (F_(1,22)_=16.171 p=0.0006). The CC04 and CC41 strains were behaviorally divergent, as previously published, with CC04 male and female mice responding to cocaine and the CC41 mice being unresponsive to the drug (**Figure 4)**. One mouse was excluded as a behavioral outlier (male, CC04, 25760 ABX). Pairwise t-tests at each time point show no significant difference between treated and control mice. However, a repeated measures ANOVA on each sex of mice showed that in females there was a time x strain x treatment effect [F_(7,13)_ = 1.692784 = p = 0.0356] Post-hoc contrast analysis showed CC04 antibiotics vs control F_(7,13)_= 3.64, p= 0.0156; whereas CC41 antibiotics vs control F_(7,13)_=0.01318, p = 1. Univariate effect of strain F_(1,10)_ = 70.874, p <0.001. However in males, there was a univariate effect of strain F_(1,18)_ = 38.4406, p<0.001 but not a significant time × strain × treatment F=_(7,12)_ = 0.630, p = 0.4407.

**Figure 3.**
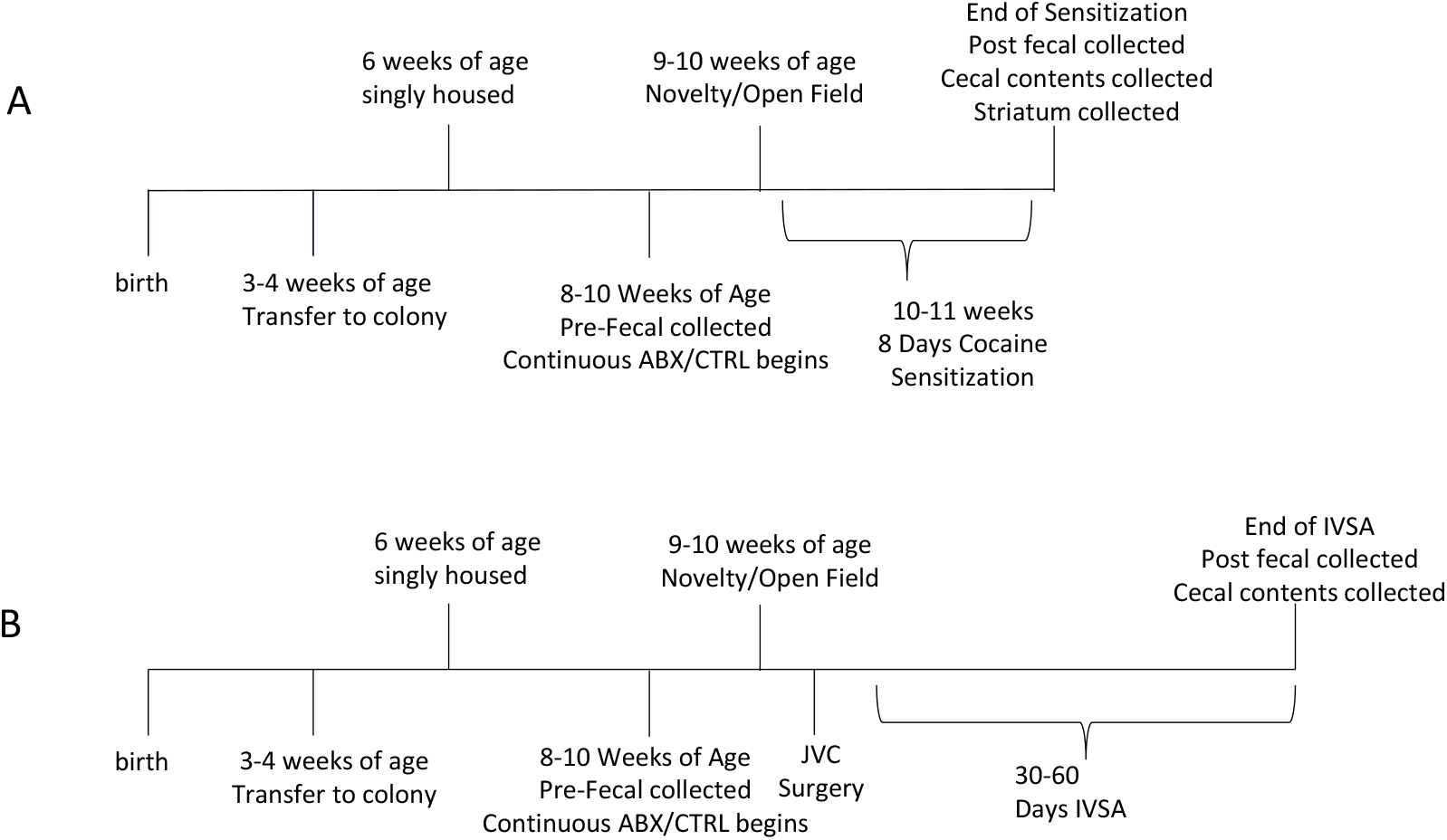
Schematic diagram of the experimental design for Antibiotic ablation of the microbiome of CC04 and CC41 mice. **A**. cocaine sensitization with 5mg/mg and **B**. cocaine intravenous self-administration. A repeated measures ANOVA on each sex of mice showed that in females there was a time x strain x treatment effect [F_(7,13)_ = 1.692784 = p = 0.0356], driven by the differences observed in CC04.

**Figure 4.**
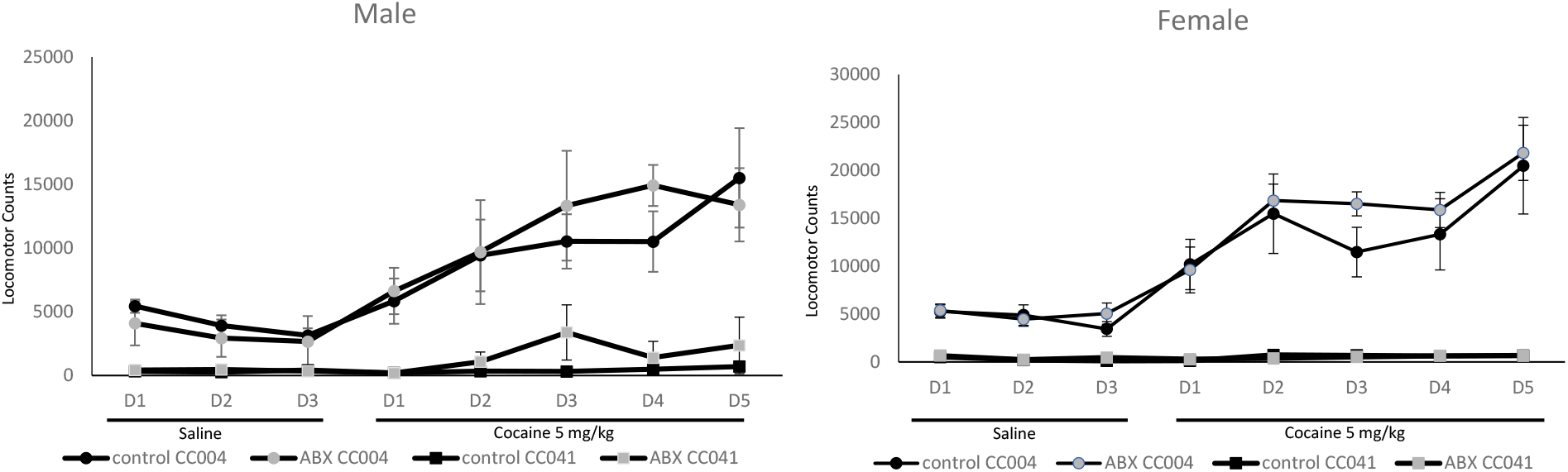
Cocaine behavioral sensitization of male and female CC04 and CC041 mice in response to antibiotic treatment and with control. **A**. The experimental timeline for the behavioral sensitization of Kiraly [12] which included three days of saline treatment followed by five days of 5 mg/kg i.p. cocaine administration. **B**. The experimental timeline for the cocaine intravenous self-administration experiment which included phases of acquisition, stabilization, and maintenance (dose response).

### 3.4 Effect of antibiotic treatment on the microbiome of CC strains that vary in response to cocaine

16S and mWGS were performed on the **cecal** contents on all mice at the end of the cocaine sensitization paradigm. ABX treatment significantly reduced the bacterial abundance in both the CC04 and CC41 strains as seen in WGS read counts (**Figure S3**). Consistent with fecal microbiome data, CC04 showed a higher abundance of *Barnsiella* (W=30, Z=-2.85, p=0.004, q=0.04, r=.86) post sensitization in comparison to CC41 post sensitization in the cecal microbiome. CC41 showed a higher abundance of *Eisenbergiella* (W=3, Z=-2.15, p=0.03, q=0.1) compared to CC04 post sensitization (**Figure 5, Table S4**). No comparisons were made between ABX samples as the reads were too low due to the efficacy of the antibiotic ablation. mWGS of cecal contents obtained from mice post cocaine sensitization showed an abundance of *Lactobacillus johnsonii* (W=30, Z=-2.85 p=0.004, q=0.26, r=0.86) and *Muribaculum intestinale* (W=30, Z=-2.83, p=0.004, q=0.26, r=0.86) in CC04. *Akkermansia muciniphila* (W=3, Z=-2.166, p=0.03, q=0.48, r=.65) were dominant in CC41 mice. *Duncaniella* abundance was also different (W=30, Z=-2.85, p=-0.004, q=0.26) with higher levels found in CC04 (**Figure 5B, Table S4)**. Different results obtained by 16S versus mWGS data profiling are to be expected based upon the type of reference database available for each technique. To produce more comparable data, 16S data was mapped to the PathSeq database used for analyzing WGS data. In this analysis *Lactobacillus* and *Akkermansia* trends remained the same (**Figure S4**).

**Figure 5.**
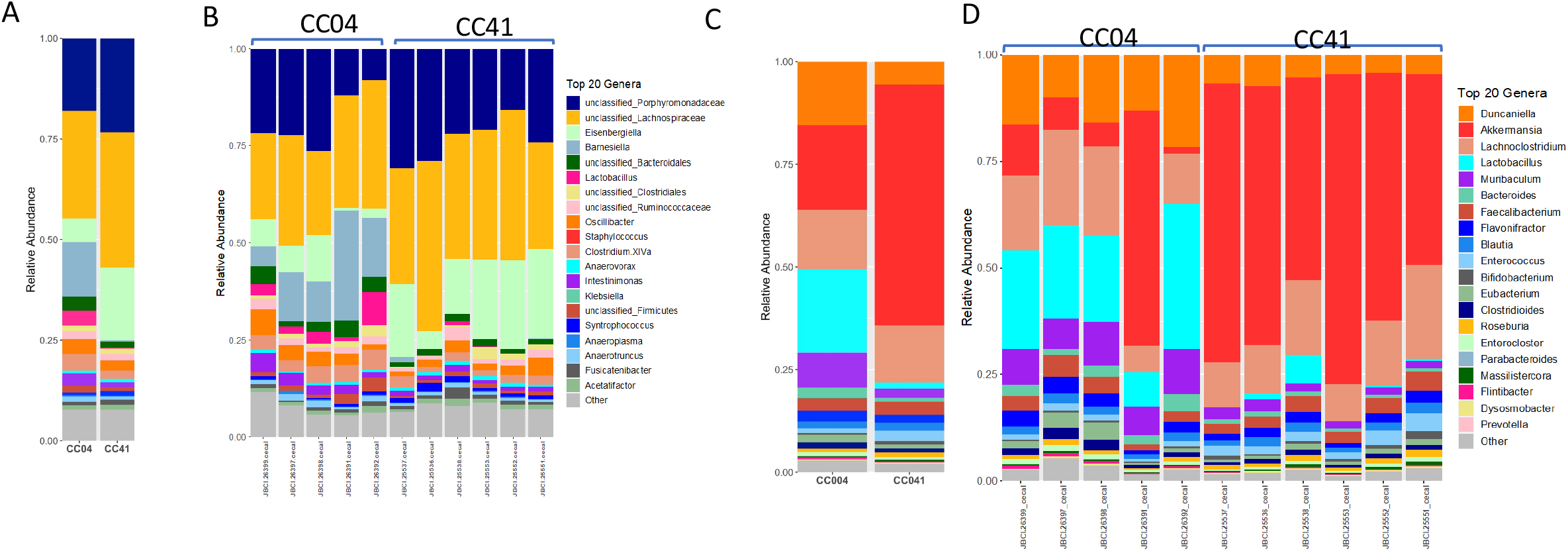
Microbiome composition of CC04 and CC41 cecal contents by 16S and Whole Genome Shotgun sequencing of control mice following cocaine sensitization. Differences in *Lactobacillus* and *Akkermansia* trend similarly in both methods of microbiome analysis. **A**. Strain average microbiome composition of the top 20 genera in cecum by 16S. **B**. Individual mouse level microbiome composition of the top 20 genera in cecum by 16S. **C**. Strain average microbiome composition of the top 20 genera in cecum by whole-genome shotgun sequencing. **D**. Individual mouse level microbiome composition of the top 20 genera in cecum by whole-genome shotgun sequencing.

### 3.5 PICRUSt analysis of the cecal microbiome of 16S data identifies differential KO modules between CC04 and CC41 strains after cocaine treatment

We have previously observed KO cluster differences in CC04 in response to cocaine that were absent in the CC41 strain. Based upon PICRUSt analysis using control samples, 68 KO modules were significantly different across the CTRL strains cocaine-sensitization treatment (**Figure 6A,B Table S5)**. Of those KO modules 20 were associated with GABA/glutamine/glutamate metabolism and seven related to short-chain fatty acid metabolism (acetate, butyrate, propionate). The KO modules K11102 other_glutamate and K13923 Propionate synthesis had the largest fold change (LFC -5.1) between the strains.

**Figure 6.**
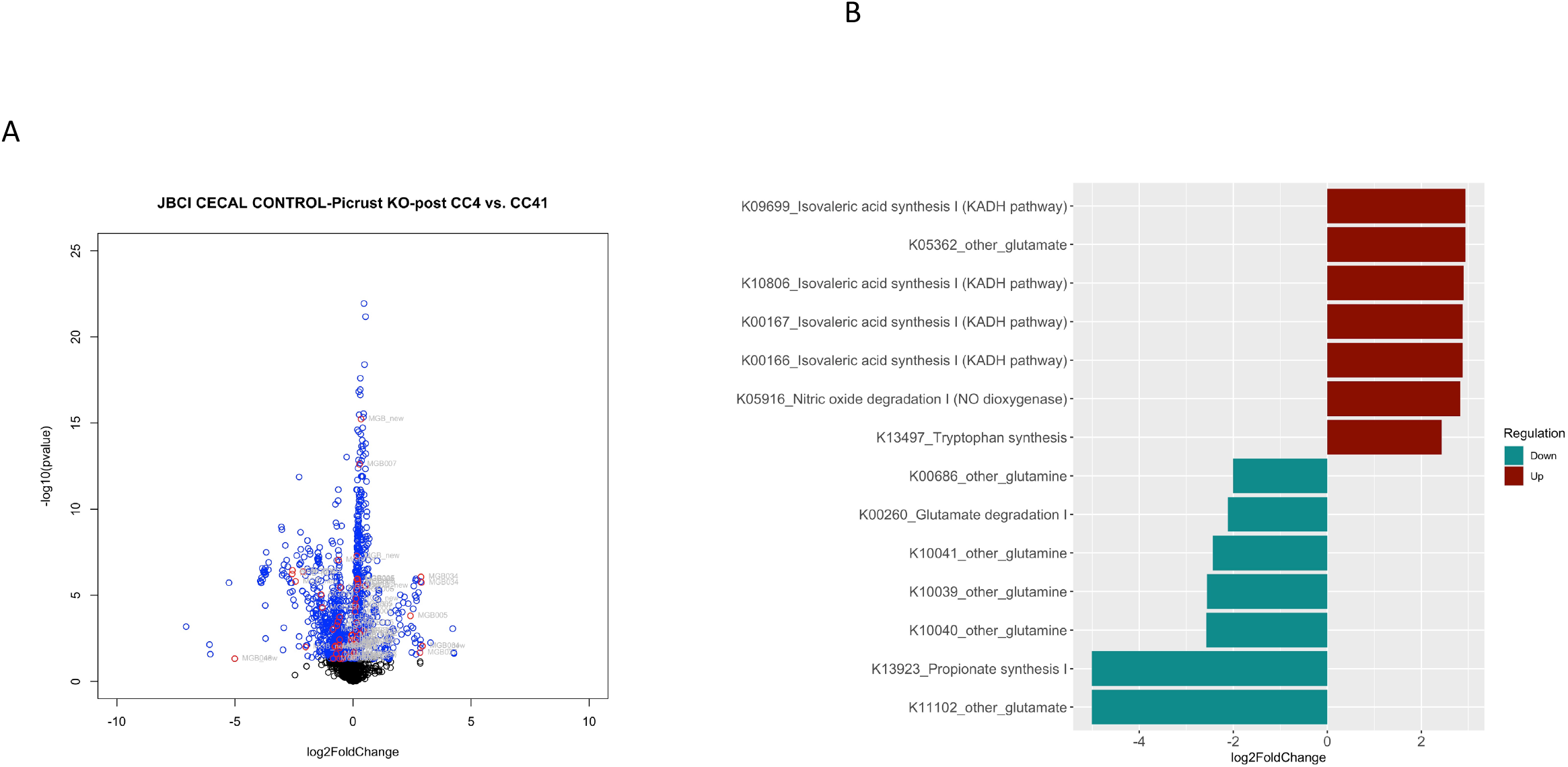
Volcano plots of the functional categories encoded by the microbiome 16S PICRUSt and the modules that are differential between strains after cocaine sensitization. **A**. A scatterplot showing the statistical significance and magnitude of change of the KO clusters as determined by PICRUSt2 of 16S data. **B**. The gut-brain module KO categories (qed<0.05) from PICRUSt2 of 16S data

### 3.6 Pathway analysis using mWGS cecum data

Utilizing the mWGS samples, we looked at the representation of the GBM pathways in the control CC04 vs CC41 samples post cocaine sensitization protocol. 13 pathways were down regulated and 14 up regulated in CC04 compared to CC41 mice (**Figure 7 and Table S6**). PWY.4321.L.glutamate.degradation.IV (LFD -3.317, p=0.0001, q=0.001) was more abundant in CC04, as was PWY.5044 purine nucleotide degradation (LFC=1.78, p=0.006, q=0.025) and PWY6666.2.dopamine.degradation (LFC -4.943, p=0.016, q=0.05).

**Figure 7.**
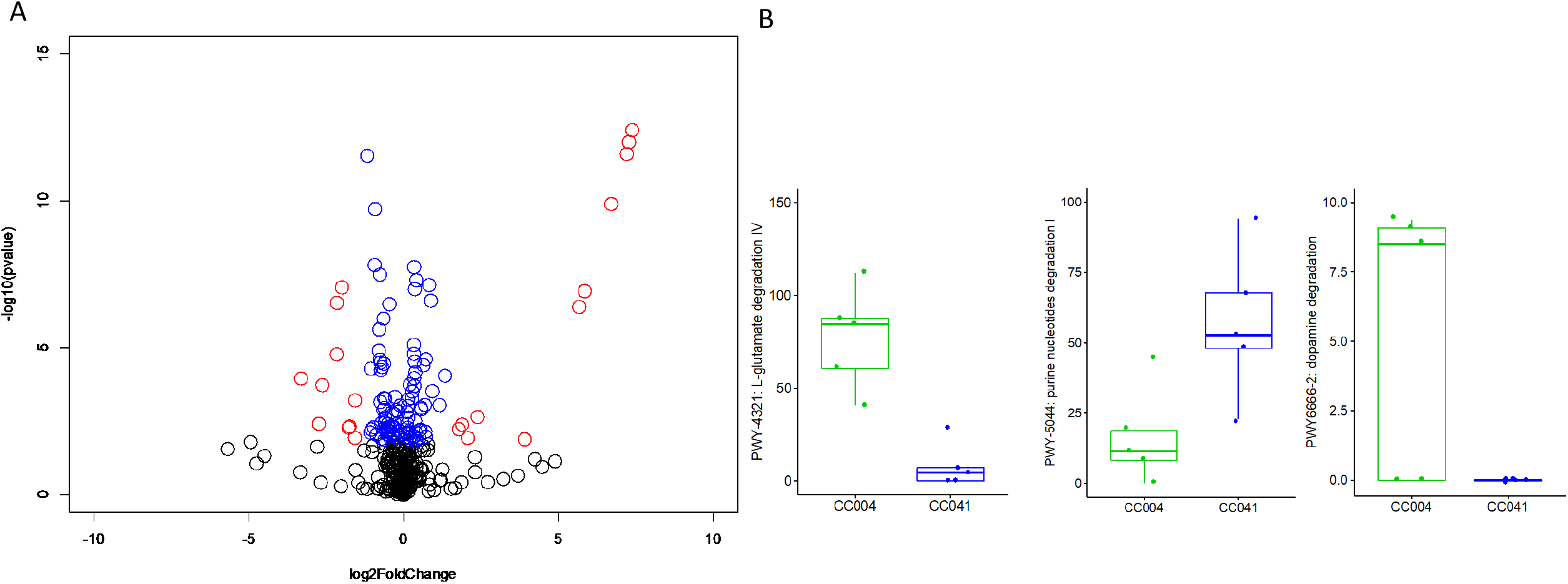
The WGS pathways that are differential in the cecal feces between CC04 and CC41 post sensitization. **A**. A scatterplot showing the statistical significance and magnitude of change of the KO pathways from WGS sequencing. **B**. Example pathways differential representation across CC04 and CC41 mice after cocaine sensitization.) Blue=significant KO/pathway (p adjusted value<0.05), red=significant KOs that are GBM (p adjusted value<0.05).

### 3.7 Antibiotic treatment effects on striatal neurotransmitter levels

We observed a significant difference in DOPAC (p<0.0001), HVA (p<0.001), 3-MT (p=0.0077) and DA (p=0.008) between CC004 and CC0041 strains before antibiotic treatment. There was no significant effect of strain by treatment, except for DA levels. For DA, the full model there was a strain × treatment effect F_(3,41)_=4.4089, p=0.0093. In a post hoc contrast of the control group there was a significant difference between CC04 vs CC041. However in the antibiotic group there was not a significant difference F_(1,38)_=4.0438, p=0.0515. In summary, strain differences in striatal dopamine were diminished by antibiotic treatment (**Table 1**).

**Table 1-.** The mean and standard error of various neurotransmitters in the striatum of CC04 and CC041 mice with out without antibiotic treatment. NE=Norepinephrine, DOPAC=3,4-Dihydroxyphenylacetic acid, DA=dopamine, 5-HIAA= 5-hydroxyindoleacetic acid, HVA= homovanillic acid, 5-HT=serotonin, 3-MT= 3-methoxytyramine. CTRL=Control treatment water with aspartame sweetener. ABX=triple antibiotic cocktail with aspartame sweetener. Measurements are reported as ng/mg total protein.

| Strain | Treatment | NE | DOPAC | DA | 5-HIAA | HVA | 5 HT | 3 HT |
| --- | --- | --- | --- | --- | --- | --- | --- | --- |
| CC004/TauUncJ n=10 | CTRL | 4.1 ± 0.6 | 7.7 ± 0.5 | 164.3 ± 8.2 | 4.8 ± 0.2 | 14.4 ± 0.7 | 16.2 ± 0.7 | 14.3 ± 0.7 |
| CC041/TauUncJ n=11 | CTRL | 3.4 ± 0.5 | 11.3 ± 0.5 | 203.9 ± 6.1 | 5.2 ± 0.3 | 17.9 ± 0.5 | 17.9 ± 1.2 | 12.2 ± 0.4 |
| CC004/TauUncJ n=11 | ABX | 4.2 ± 0.5 | 9.0 ± 0.7 | 169.8 ± 5.8 | 4.9 ± 0.2 | 13.9 ± 0.6 | 17.2 ± 1.0 | 15.0 ± 0.7 |
| CC041/TauUncJ n=10 | ABX | 3.6 ± 0.3 | 11.5 ± 1.0 | 196.2 ± 15 | 5.4 ± 0.4 | 16.8 ± 1.0 | 19.1 ± 1.1 | 13.0 ± 1.1 |

### 3.8 Antibiotic treatment effects on cocaine intravenous self-administration

The same antibiotic treatment approach used for cocaine sensitization was repeated with cocaine intravenous self-administration. This time cecal weights were collected and were determined to be significantly different between antibiotic and control treated samples (F_(1,18)=_414.25, p < 0.001). Surprisingly there was a strain × treatment effect (F_(3,21)_=190.326, p <0.001) on cecal size as well. The cecums from CC041 being significantly larger than the cecums of CC04 after antibiotic treatment. In a post-hoc contrast there no difference between the cecum weights of the strains from the control sample (F_(1,18)_=1.6645, p=0.213) (**Figure S5**).

Analysis of the IVSA dose response curve by performing pairwise t-tests at each dose for each strain showed no significant differences from treatment, at any one dose or in a particular strain (**Figure 8**). In a repeated measures MANOVA across the dose response curve there was no significant dose × strain × treatment interaction (F_(4,15)_=0.521, p<0.1513). There was a significant dose x strain interaction (F_(4,15)_=10.01, p<0.0001) and a dose × treatment interaction (F_(4,15)_=0.876, p=0.041) likely driven by the antibiotic treated CC04 dose response. This was supported by a post-hoc contrast analysis of the dose × strain × treatment interaction, including only CC04 the antibiotic and control treamanet that showed F_(4,15)_=1.34, p=0.0088.

**Figure 8.**
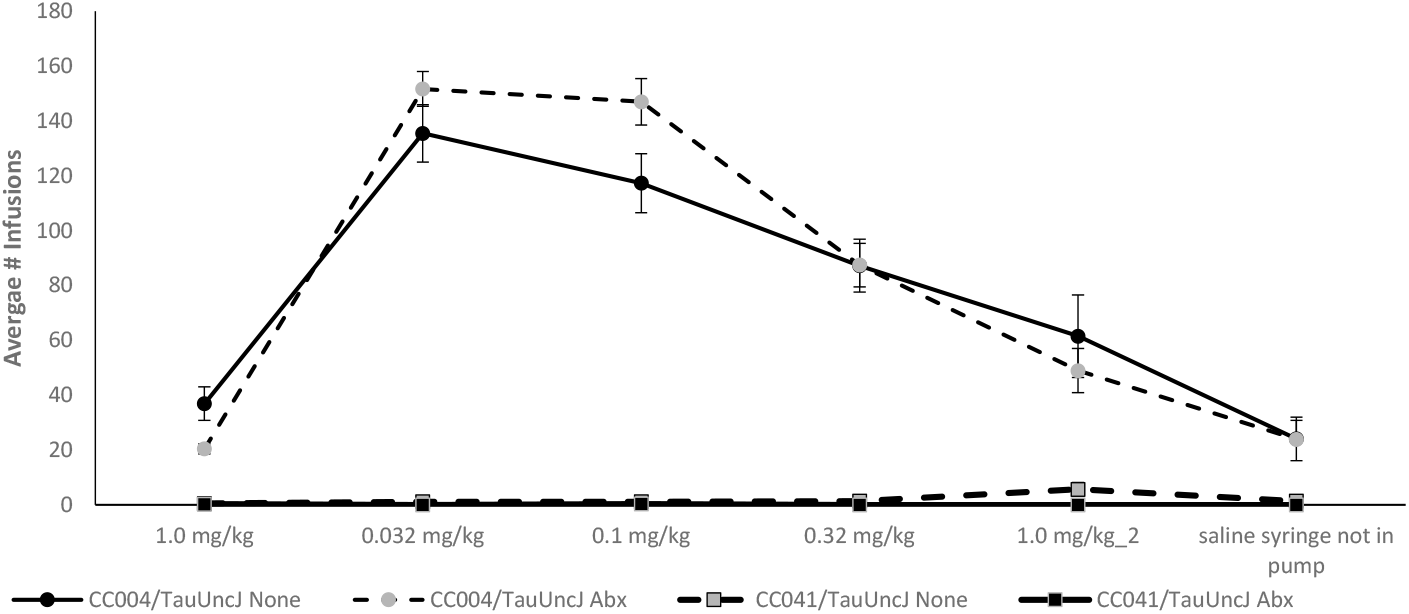
Cocaine intravenous self-administration dose response curve of male CC04 and CC041 mice in response to antibiotic treatment and or control. The mean and standard error for the total number of infusions administered at each dose for each strain and treatment was plotted against the dosage.

## 4. DISCUSSION

Extensive studies have shown that host genetics have a causal role in the structure and composition of the gut microbiome [8, 9]. We have previously identified QTL regulating the abundance of numerous bacteria in the CC breeding population [7], and demonstrate here how two behaviorally divergent CC strains [18] also have unique microbiome compositions. We have confirmed that repeated cocaine treatment of male and female CC04 mice results in an upregulation of *Barnsiella* as was reported in the colon of male C57BL/6N mice [32]. We have also shown that the effects of the cocaine sensitization paradigm upon the microbiome are strain specific and that an intact microbiome (not antibiotic treated) does not significantly repress a low dose cocaine sensitization response as observed in C57BL/6J males [12]. We did observe that naïve strain differences in striatal DOPAC between CC04 and CC41 are abolished by antibiotic treatment. These findings are consistent with host genetics controlling both the differential behavior observed in these two strains, their innate microbiome composition, and the microbiome’s response to repeated drug exposure.

CC strains have been observed to express a great deal of variation in drug-related traits[18, 33-35]. Phenotyping this reference panel of RI lines has revealed phenotypic variation much more extreme than originally observed in the eight inbred founder strains. The microbial differences include numerous taxa such as *Acetatifactor, Amerpfustis, Anaeroplasma, Butyrivibrio, Catabacter, Christensenella, Eisenbergiella, Enterococcus, Klebsiella, Lactobacillus, Parvibacter, Robinsoniella, Roseburia, Ruminococcus, Syntrophococcus* and several unclassified microbes among the 82 most abundant genera.

One of the surprising results of this study was the observation that antibiotic treatment did not dramatically increase behavioral sensitization to a dose of 5 mg/kg cocaine in male and female mice of the CC04 and CC41 strains. Instead, we observed a sex-specific response to antibiotics in CC04 female mice that was not observed in CC04 males or either sex of CC41. While the published data showed differences in cocaine sensitization behavior at 5 mg/kg, there were no behavioral differences of antibiotic treatment at 10 mg/kg in the male B6J mice tested in that same publication[12]. This suggested that the repressive effect of the microbiome on sensitization behavior may be dose dependent. In our antibiotic experiment, we tested to a single dose of cocaine (5mg/kg) and perhaps CC04, a very high responder would have required antibiotic treatment of a lower dose of cocaine to remove the repressive effect of the microbiome on sensitization behavior. Similarly, CC41 mice exhibit no locomotor response to cocaine at the published dose of 10mg/kg [18] and 5 mg/kg might have shown a more dramatic behavioral response at a higher concentration of cocaine, that could be altered by the microbiome. Our results, together with those of Kiraly suggest that the microbiome effect on locomotor sensitization in response to cocaine is dependent on strain, sex and dose of cocaine used [12].

We performed cocaine IVSA on a small cohort of CC04 and CC041 mice and detected a significant difference in response to antibiotics for strain CC04 that we did not detect with strain CC41. Although no doses individually differed between the CC04 with and without antibiotics there was a trend where at the lower doses, 0.032 mg/kg and 0.1 mg/kg there were more infusions in the antibiotic treated CC04 then the control. This supports the observation of Kiraly in the cocaine sensitization that the antibiotic effect appears to be dose dependent and here demonstrates that the effect of antibiotics is also strain dependent in cocaine IVSA.

One of the most dramatic differences is the abundance of *Eisenbergiella* in CC41, a strain that shows little locomotor response to cocaine and an increase in *Lactobacillus* in CC04 strain that is highly responsive to the locomotor effects of cocaine. It is interesting to note that the strain that is not behaviorally responsive to cocaine did not have any microbiota that were significantly different before and after cocaine administration, which is in stark comparison to CC04 which is very responsive and exhibited a large increase in *Barnesiella* and smaller increase in *unclassified_Coriobacteriaceae*. Previously a study that treated C57BL/6N male mice with 20 mg/kg of cocaine ip for 7 consecutive days identified four genera that were decreased, *Mucispirillum, Butyricicoccus, Pseudoflavonifractor* and unclassified *Ruminococcaceae* while there was an increase in *Barnesiella*, and unclassified members of *Porphyromonadaceae, Bacteriodales*, and *Proteobacteria* [32]. It must be noted that the C57BL/6N strain of mouse in comparison to the C57BL/6J strain of mice contains a mutation in *Cyfip2* gene and presents with a relatively dampened cocaine sensitization response [36]. The authors suggest that the change in microbiome composition, dysbiosis, was associated with upregulation of proinflammatory mediators including NF-κB and IL-1β. Similarly, they show that cocaine altered the gut-barrier composition of the tight-junction resulting in a leaky gut. While the authors used only one sex, one strain and one high repeated dose of cocaine, we have used multiple strains and see the dysbiosis phenotype only in the strain that is behaviorally responsive to both doses of cocaine [12].

The differential metabolic potential that was identified using either PICRUSt or mWGS pointed to both glutamate degradation and dopamine degradation enzymes encoded by the microbiome as upregulated in CC04 but not CC41 in response to cocaine. Glutamate receptors line the epithelial cells of the gut and have been show to signal through vagal afferents to regions of the brain [37, 38]. Glutamate is also the substrate for GABA production and GABA producing strains such as *Lactococcus lactis* have been shown to modulate behavior [39]. The finding that glutamate metabolism is altered in response to cocaine in responsive strains supports the glutamate hypothesis of substance use disorder [40-42], and was supported by our recent findings in the Diversity Outbred Mice [43] extending that process into the gut. These observations provide some mechanistic insight into the gut-brain axis and SUD.

## 5. CONCLUSION

In conclusion, our study indicates that the microbiome shows changes in response to cocaine, in a strain dependent manner. Strains such as CC041 that appear behaviorally non-responsive to cocaine, also contain a microbiome that does not robustly respond to cocaine. However, strain CC04 that exhibits a strong locomotor response to cocaine also displays dramatic changes with its microbiome composition in response to both a high and low dose of cocaine. These changes in response to cocaine are detectable at the level of changes to the metabolic potential of the microbiome. CC04 has an increase in microbes with genomes containing an abundance of glutamate and dopamine metabolic pathways. Antibiotic ablation of the microbiome does not alter the sensitization response of either strain to 5 mg/kg cocaine - CC41 mice remain non-responsive and CC04 females showed only a minimal increase in locomotor activity in response to antibiotics. Finally antibiotic ablation of the microbiome altered the IVSA dose response curve in the CC04 male strain but not in the male CC41 strain.

## SUPPORTING INFORATION

**Table S1 The variable names and all the metadata generated by open field**

**Table S2-** Tab 1-Differences between CC04 and CC41.Tab 2- Differences between CC04 in response to Cocaine. Tab 3- Differences between CC41 in response to Cocaine. Pink cells represent p<0.05.

**Table S3-** Tab 1- Gut-Brain Modules from PICRUSt that differed (p<0.05) between CC04 and CC41 Tab 2- Gut-Brain Modules from PICRUSt that differed (p<0.05) between CC04 naïve and CC04 post Tarantino Cocaine Sensitization.

**Table S4-** Tab 1 Differences between cecum of CC04 and CC41 post cocaine sensitization paradigm of Kiraly. Tab 2 WGS Differences between cecum of CC04 and CC41 post cocaine sensitization paradigm of Kiraly.

**Table S5** Gut-Brain Modules from PICRUSt of 16S data that differed (p<0.05) between CC04 and CC41 in cecum samples.

**Table S6** KEGG Orthology pathways that were differentially abundant in the cecum of CC04 vs CC41 mice based upon the functional potential of the WGS data.

**Figure S1 Distance traveled over 60 minute period post dosing on the 10 mg/kg cocaine sensitization protocol**. n=5 CC04 and n=6 CC41.

**Figure S2 Weight of animals over the eight days of the cocaine sensitization protocol**. Based upon weight there was no adverse effect of the antibiotic treatment upon animal weight.

**Figure S3 Total read counts from the control and antibiotic treated mice**. There were very few reads recovered from the antibiotic treated mice, which displayed enlarged cecums consistent with a severe microbiome disruption.

**Figure S4. Microbiome composition of CC04 and CC41 cecal contents by 16S and Whole Genome Shotgun sequencing of control mice following cocaine sensitization** For this analysis the 16S data was mapped to Pathseq complete genome (90% similarity).

**Figure S5 Response of the cecum to antibiotic treatment in CC04 and CC041 mice**. (A) Cecal weights and (B) Representative image of CC41 cecum post IVSA treatment.

## ACKNOWLEDGEMENT

The authors gratefully acknowledge drug reagents from the NIDA Drug Supply Program. The Vanderbilt Neurochemistry Core is supported by the Vanderbilt University Brain Institute.

## DATA AVAILABILITY STATEMENT

The 16S and WGS data are available on SRA under accession SUB12204043.

