## Supplementary figures and images for "The microbial community dynamics of cocaine sensitization in two behaviorally divergent strains of collaborative cross mice"

### Supplemental Table 6

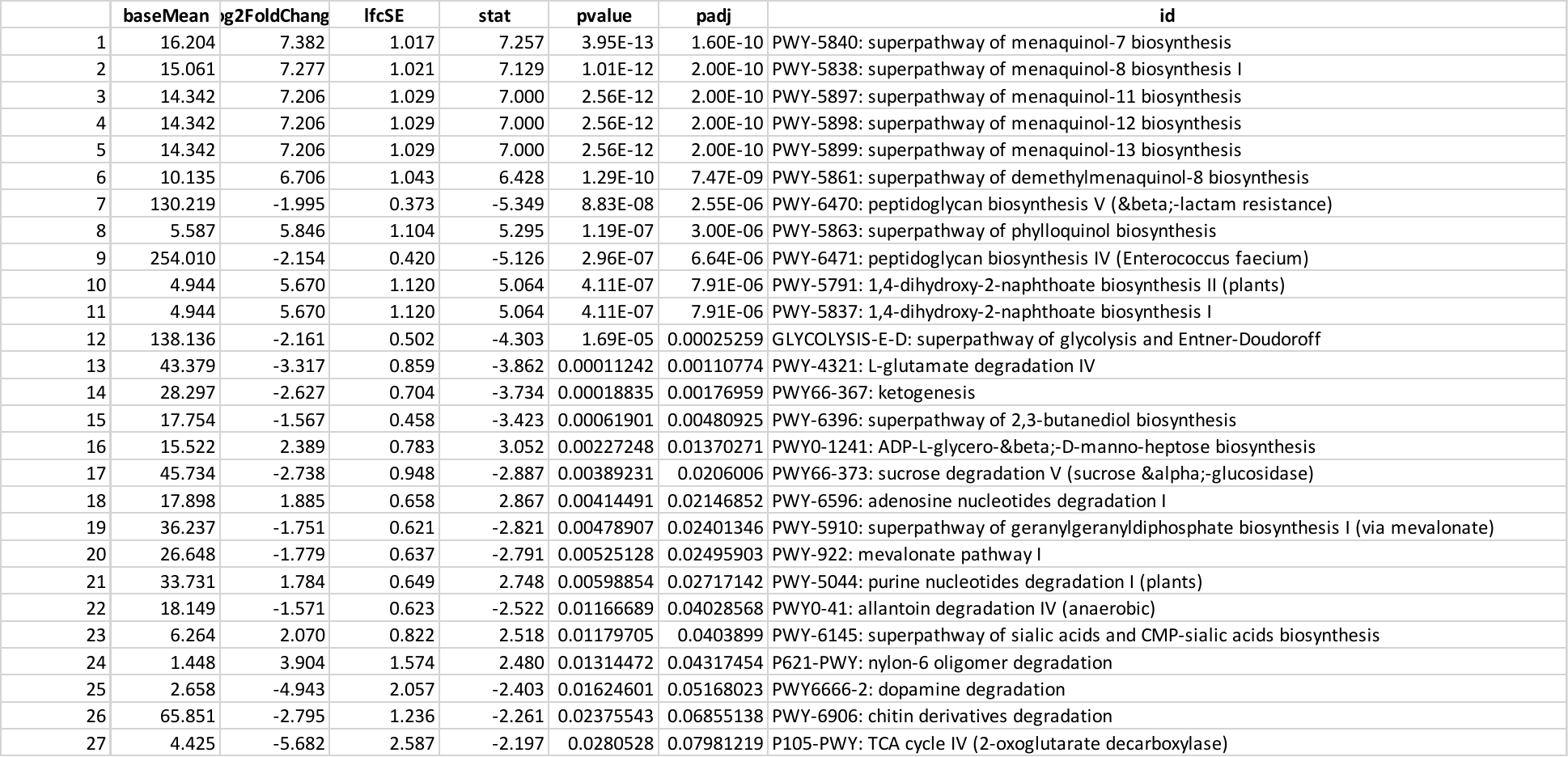
